# Single and paired TMS pulses engage spatially distinct corticomotor representations in human pericentral cortex

**DOI:** 10.1101/2024.10.03.616450

**Authors:** Mads A.J. Madsen, Lasse Christiansen, Chloe Chung, Morten G. Jønsson, Hartwig R. Siebner

## Abstract

Single-pulse transcranial magnetic stimulation (TMS) of the primary motor hand area (M1-HAND) can assess corticomotor function in humans by evoking motor evoked potentials (MEP). Paired-pulse TMS at peri-threshold intensity elicits short-latency intracortical facilitation (SICF) with early peaks at inter-pulse intervals of 1.0-1.8ms (SICF_1_) and 2.4-3ms (SICF_2_). The similarity between the periodicity of SICF and indirect (I-)waves in the corticospinal volleys evoked by single-pulse TMS suggests that SICF originates from I-wave generating circuits. This study aimed to explore the mechanisms of MEP generation by mapping the corticomotor representations of single-pulse and paired-pulse TMS targeting SICF_1_ and SICF_2_ peaks in 14 participants (7 female). MEPs were recorded from two hand muscles and the spatial properties of each corticomotor map were analyzed. For both hand muscles, we found a consistent posterior shift of the center-of-gravity (CoG) for SICF maps compared to single-pulse maps, with a larger shift for SICF_1_. CoG displacement in the SICF_1_ map correlated with individual SICF_1_ latencies. Further, ADM maps consistently peaked more medially than FDI maps and paired-pulse TMS resulted in larger corticomotor maps than single-pulse TMS. This is the first study to show that circuits responsible for SICF have a more posterior representation in the precentral crown than those generating MEPs via single-pulse TMS. These findings indicate that paired-pulse TMS probing SICF_1_, SICF_2_, and single-pulse TMS engage overlapping but spatially distinct cortical circuits, adding further insights into the intricate organization of the human motor hand area.

**New & Noteworthy:** Single- and paired-pulse transcranial magnetic stimulation (TMS) is widely used to study corticomotor physiology in humans, but do they engage the same intracortical circuits? We compared the spatial properties of corticomotor maps elicited by single-pulse TMS to those elicited by paired-pulse short-latency intracortical facilitation (SICF). SICF maps consistently showed a posterior shift in center of gravity compared to single-pulse maps, suggesting that paired-pulse TMS engages cortical circuits that are spatially distinct from single-pulse TMS.

## Introduction

Transcranial magnetic stimulation (TMS) of the hand motor representation in the precentral gyrus (M1_HAND_) can cause a contraction of contralateral hand muscles, making TMS a powerful non-invasive tool to probe corticomotor physiology. Exactly which cortical elements are stimulated by TMS is still unknown, but biophysical modelling of morphologically realistic cortical neurons suggests that myelinated axon terminals from pyramidal cells and inhibitory interneurons in the crown of the gyri constitute low-threshold targets for TMS (1-3). Interestingly, anatomical tracer studies in non-human primates show that the primary output region of monosynaptic pyramidal tract neurons (PTNs) in the primary motor cortex lies in the evolutionary younger, posterior part of Broadmann’s area 4, which is typically located in the sulcal wall or fold (4). Additionally, human epidural recordings show that TMS elicits both a direct descending wave followed by a number of later indirect (I)-waves (5) suggesting that populations of PTNs are activated trans-synaptically and experience numerous discharges from a single TMS pulse. The notion is supported by paired pulse TMS protocols where an additional TMS pulse (usually sub-threshold) is used to condition the response to the single TMS pulse alone, often resulting in marked changes in peak-to-peak amplitude of the motor evoked potential (MEP)(6, 7). For example, two TMS pulses delivered in very close temporal proximity i.e. interstimulus intervals (ISIs) of ∼1-5ms markedly facilitates the MEP as compared to single-pulse TMS (8). This phenomenon, coined short-latency intracortical facilitation (SICF)(9), is cortically mediated (9, 10) and more pronounced when interstimulus intervals mimic the periodicity of the I-waves observable in epidural recordings, although also detectable at SICF trough ISIs with higher stimulation intensity or more pulses (11).

Because the periodicity of SICF matches that of epidurally recorded I-waves the general hypothesis is that SICF works through facilitation of the I-wave generating circuits that are also involved in generating single-pulse MEPs. However, the exact cortical networks involved in generating single-pulse MEPs and SICF and their similarities are still unknown. Moreover, several findings suggest that the first and second SICF peaks (SICF_1_ and SICF_2_) may reflect activity in non-identical cortical circuits that subserve distinct sensorimotor functions (12-14). While substantial effort has been made to pinpoint the most likely site of cortical activation by single-pulse TMS (3), the spatial location of the intraneuronal networks underpinning SICF remains entirely unexplored.

The objective of this study was to map the spatial cortical representation of single pulse TMS and SICF in order to test whether spatially distinct or identical cortical populations underpin these phenomena. Our hypothesis was that paired pulse SICF engages different neural populations, compared to single pulse, evident as a systematic difference in the center of gravity of the corticomotor maps. Additionally, we hypothesized that there would be a spatiotemporal relationship between the spatial displacement and the ISIs of the first and the second SICF peak. To test these hypotheses, we mapped the peri-central corticomotor representations of intrinsic hand muscles with single-pulse TMS and two SICF conditions with ISIs adjusted to the individual SICF_1_ and SICF_2_ peak latency respectively. We used central sulcus shape-based mapping (15, 16) as it provides higher sensitivity towards muscular somatotopy compared to standard grid-based mapping (16, 17) and is sensitive towards between subject variability in hotspot location (18).

## Materials and Methods

### Experimental design

Before the experimental TMS session, participants underwent a structural T1-weigthed magnetic imaging resonance (MRI) scan, which was used to plan TMS targets for sulcus-shaped mapping (16). In the experimental session participants went through recordings of SICF curves to determine individual latencies of SICF peaks. Following this, three different robot-assisted, neuro-navigated, sulcus shaped TMS motor maps were created using single- or paired-pulse TMS targeting either the first or the second SICF peak (Fig. 1).

**Figure 1.**
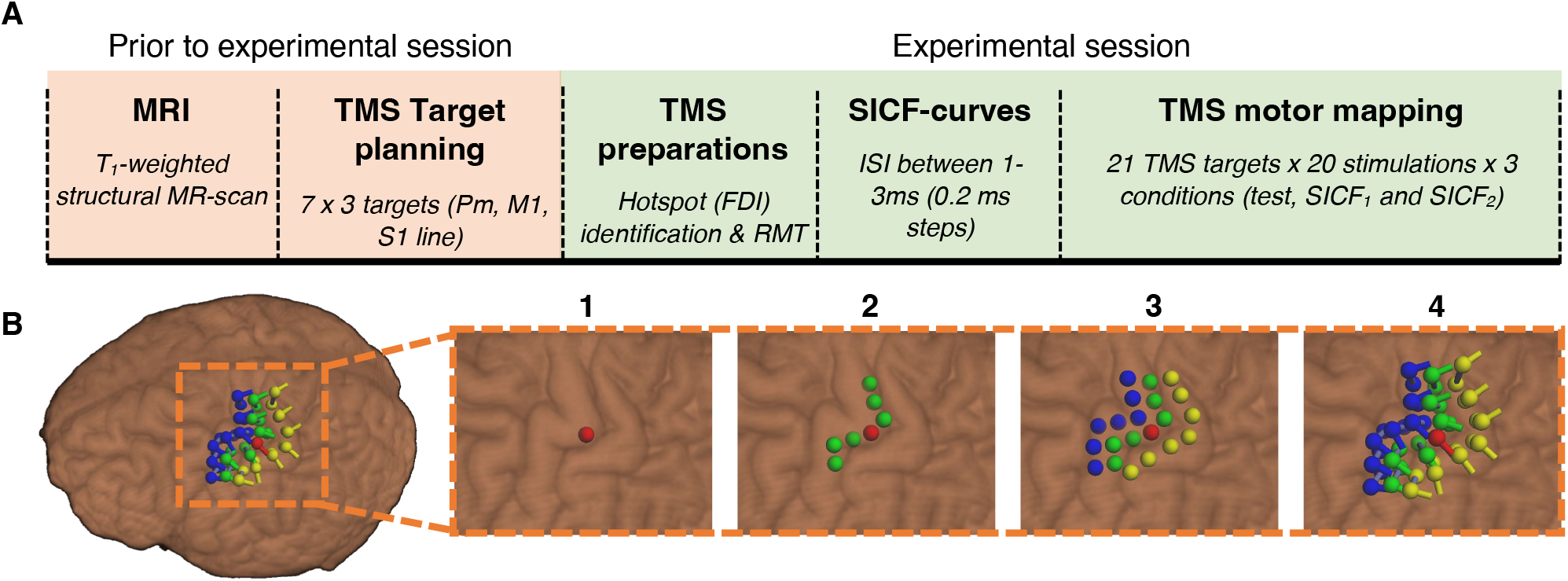
Experimental design and protocol. **A**) Flow chart of the experimental procedure. **B**) Map planning in the neuronavigation software for the robot assisted TMS motor mapping. Abbreviations: MRI = magnetic resonance imaging, TMS = transcranial magnetic stimulation, FDI = first dorsal interosseous, SICF = short intracortical facilitation, Pm = premotor cortex, M1 = primary motor cortex, S1 = primary sensory cortex.

### Participants

Seventeen healthy right-handed volunteers (mean age: 26.9 years, 8 females) participated in the study. Handedness was assessed by the Edinburgh handedness inventory (19). All participants had no history of psychiatric disorders and were screened for contraindications to TMS (20). Written informed consent was obtained from all subjects and studies were performed in accordance with the Helsinki declaration on Human experimentation. The study was approved by the Ethics Committees of the Capital Region of Denmark (H-15000551). Two participants were excluded from the study due to continuous background EMG contractions and one participant was excluded due to a too high resting motor threshold (>91%MSO). Thus, 14 subjects (7 females) were included in the statistical analyses.

### Magnetic resonance imaging

Prior to the experimental session, a three-dimensional T1-weighted magnetization-prepared rapid gradient-echo (MPRAGE) magnetic resonance (MRI) scan was acquired on a 3T Verio scanner, (Siemens, Erlangen, Germany) with the following parameters: TR/TE=2300/2.98ms, T1=1100ms; 256 × 256 matrix, flip angle 9°, 1mm^3^ isotropic voxel).

### Surface electromyography

Electrical muscle activity of the right first dorsal interosseus (FDI) and abductor digiti minimi (ADM) muscles were recorded with surface electrodes (Ambu Neuroline 700, Ballerup, Denmark) using a bipolar belly-tendon montage. The Analog EMG signal was filtered (band-pass, 5-2000 Hz), amplified (x1000, Digitimer, Cambridge Electronic Design (CED), Cambridge, UK), digitalized (sampling frequency 5000 Hz, 1201 micro Mk-II CED), recorded (Signal 4.0, CED), and stored on a computer for later off-line analysis. During the experiment, background electromyography (EMG) was monitored by the experimenter, and participants were instructed to relax whenever background EMG was detected.

### Transcranial Magnetic Stimulation

Single- and paired-pulse TMS was delivered using a MagPro x100 Option stimulator (Magventure, Skovlunde, Denmark) connected to a cooled-MC-B35 figure-of-eight coil with windings of 35mm diameter (Magventure). A biphasic pulse configuration that induced an antero-posterior followed by postero-anterior (AP-PA) current in the cortex was selected to evoke MEPs at the lowest possible stimulation intensity (21). All TMS was performed using stereotactic neuronaviation (Localite, Sankt Augustin, Germany) in which individual structural MRI-images were uploaded. The optimal coil position (hotspot) to activate the right FDI was identified through a mini-mapping procedure and defined as the site where TMS elicited the largest and most consistent MEPs. Resting motor threshold (RMT) was determined to the nearest 1% of maximum stimulator output using single-pulse TMS and defined as the minimal stimulus intensity required to evoke a response of 50µV in at least 5 out of 10 successive trials (22). Following the determination of the hotspot and RMT, TMS was performed by a neuronavigation TMS robot (Axilum Robotics, Schiltigheim, France) for the remainder of the experiment.

### Short-interval intracortical facilitation (SICF) curves

For paired-pulse TMS to induce SICF, the intensity of the first test stimulus (TS) was set to 110% RMT (determined for FDI) and to 90% RMT for the second conditioning stimulus (CS)(8). An SICF-curve was created before the mapping procedure to determine individual SICF_1_- and SICF_2_ -peak latencies. Inter-stimulus intervals (ISI) ranging from 1.0ms to 3ms were tested in 0.2ms steps. Ten paired TMS pulses were delivered in pseudo-random order over the FDI hotspot for each ISI together with test stimuli alone. The average peak-to-peak MEP amplitude was calculated online for each condition and plotted against the respective ISI. The intervals that produced the largest average peak-to-peak MEP amplitude for the FDI muscle, within the first (1-1.8ms) and second (2.4-3ms) SICF range were determined as individual SICF_1_ - and SICF_2_-peak latencies.

### Robot assisted, neuro-navigated, sulcus-shaped TMS motor mapping

#### TMS mapping targets

TMS mapping was done in a 3×7 grid centered around the hand knob of the precentral gurus (23) which was identified by a trained investigator. Seven targets were placed along the posterior part of the crown of the precentral gyrus (i.e. the gyral ‘lip’)(see Fig. 1B). Each target was placed conforming with individual shape of the central sulcus to form a line (M1-line). The posterior convexity of the “hand-knob” was used as a central reference point which corresponded to target 4. Coil orientation was adjusted to produce a current direction perpendicular to the central sulcus, which has been shown to be superior for M1-mapping (16). Two additional lines were created to examine the potential antero-posterior shift of somatotopy with SICF mapping. A premotor (Pm) line was positioned in accordance with the M1 line but with an anterior shift such that the targets were placed on the precentral gyral crown. Additionally, seven posterior targets were placed in accordance with the M1-line but on the crown of the postcentral gyrus (S1-line).

#### TMS motor mapping procedure

Three motor maps were conducted in the same experimental session. For each map 20 single (test condition) or paired (SICF_1_ and SICF_2_ condition) TMS pulses were applied to each of the 21 targets in a random order. The order of the three conditions was randomized and counter-balanced across subjects and TMS pulses were delivered at a frequency of 0.25 Hz (25% jitter). SICF and single pulse TMS maps were generated separately rather than intermixed to avoid the increased MEP amplitudes from SICF to potentially influence single pulse MEP amplitudes (24).

### Data analyses

#### EMG

EMG data was processed in Matlab (version R2017b, The MathWorks Inc., Massachusetts, USA) and R-studio (R-core team, 2021). All EMG traces were visually inspected and discarded if any background muscle contraction was detected. Peak-to-peak amplitude of the MEP was calculated from the remaining trials in the interval 15-40ms after the TMS pulse. The average MEP amplitude was calculated for each individual and within each muscle for either each ISI (SICF-curves) or for each stimulation position and condition (motor maps). Any target with a mean MEP amplitude <0.05mV was interpreted as a non-response site and the MEP value set to 0 to exclude the contribution of background noise to the results.

#### TMS motor maps

##### Facilitatory effects of SICF mapping

The corticomotor representation of SICF was assessed as the area and volume of the generated maps. The area of the motor maps was computed as the number of TMS targets eliciting an average MEP response of above 50µV. Additionally, map volume was calculated as the sum of the mean MEP amplitude from each target position within each motor map. Map volume was calculated both from raw MEP amplitudes and normalized to the largest mean MEP of each map (i.e., “peak-normalized” volumes).

#### Center of gravity

TMS target positions were extracted from the neuronavigation software as 3-D coordinates and converted into MNI space using SPM affine matrix conformation. As the hand-knob is not aligned to any one axes in 3-D cartesian space and because the grid of TMS targets closely resembled a plane, we reduced the dimensionality of the maps, by performing a principal component analysis (PCA) on centered target-coordinates of each individual map. The resulting first two principal components were used to construct 2-D maps where the first principal component (PC1) primarily represented the medio-lateral but also the dorso-ventral axis (referred to as the medio-lateral axis) and the second principal component (PC2) primarily represented the anterior-posterior axis. From the 2-D maps we calculated the amplitude weighted mean position (i.e. the center of gravity (CoG)) for both principal components, using the following formula:

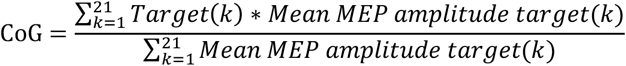

Where target(k) refers to the PC1 or PC2 coordinate for each of the 21 TMS target positions and mean MEP amplitude target(k) refers to the mean peak-to-peak MEP amplitude at each target. The CoG was computed for each condition (test, SICF_1_ and SICF_2_) and for each muscle (FDI or ADM) separately. Along with CoG coordinates we also calculated the Euclidian distance between FDI and ADM CoGs for all three conditions, and within the FDI and ADM muscle between test and SICF_1_ and SICF_2_.

### Statistical analysis

Statistical analyses were carried out in R (R core team, 2021). For all analyses, linear mixed effect models were fitted using the *lme4* and *lmertest* packages (25, 26), and multiple comparisons and adjusted p-values were computed using the *multcomp* package (27). Post-hoc pairwise comparisons were corrected for multiple comparisons using the ‘single-step’ method (27) unless stated otherwise and statistical significance was assumed if P<0.05. Homoskedasticity and normality was assured by visual inspection of residual- and quantile-quantile plots respectively. Log-transformation was applied to the dependent variable if any obvious deviations from these assumptions were detected. For SICF curves mean MEP amplitude was entered as the dependent variable with the interaction between MUSCLE and ISI as fixed effects and SUBJECT as a random effect with random intercept and a random slope of the effect of MUSCLE. P-values were obtained from analysis of variance tables for the fixed effects on the full model using Satterhwaite’s degrees of freedom method.

The same approach was taken when testing for differences in the CoG of the PC1 and PC2 coordinates and differences in map volume and area. Here MUSCLE and CONDITION (test, I1 and I2) were entered as fixed effects and SUBJECT as a random effect with random intercept and a random slope of the effect of MUSCLE. Differences in Euclidian distance between or within muscle were assessed with a linear mixed model with CONDITION as a fixed effect and SUBJECT as a random effect with random intercept.

## Results

### SICF curves

Biphasic paired pulse TMS produced distinct SICF peaks around the two expected ISI intervals in all participants. Analysis of log-transformed MEP amplitudes from the SICF curves for the entire cohort showed no interaction between MUSCLE and ISI (F_(11)_=0.36, P=0.97). There was however a main effect of MUSCLE, with overall larger MEP amplitudes in the FDI muscle (F_(1)_=6.06, P=0.027). There was also an ISI dependent facilitation in MEP amplitude (F_(11)_=61.4, P<0.001). In accordance with previous work using monophasic or biphasic stimuli (11, 28), the post-hoc pairwise comparisons showed that paired-pulse TMS at ISIs ranging from 1.0-1.8 ms and from 2.4-3.0 ms increased mean MEP amplitude compared to MEPs evoked by a single TMS pulse (all P<0.001)(Fig. 2A). The mode of SICF_1_ peak latency was 1.4 ms (10 subjects) and 2.8 ms for SICF_2_ latency (eight subjects) (Fig. 2B). Mean relative facilitation for the SICF_1_-peak was 445% (range: 174-827%) for ADM and 418% (range: 149-1008%) for FDI. SICF_2_-peak MEP amplitudes were 506% (range: 235-983%) relative to test for ADM and 414% (range: 176-700%) for FDI (Fig. 2C). There were no significant differences in relative peak facilitation between SICF-peaks or muscles (P>0.05).

**Figure 2.**
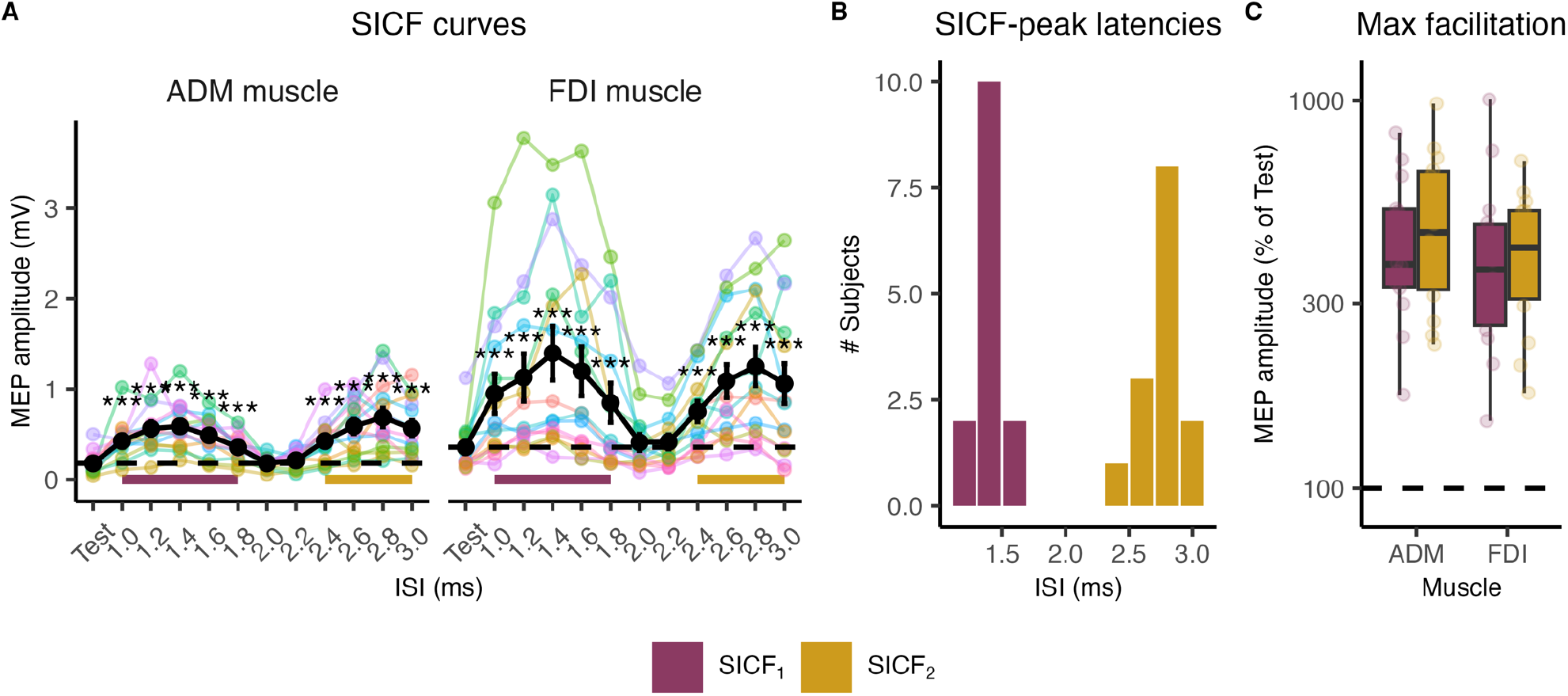
SICF curves and individual SICF-peak latencies. **A**) Individual (color) and mean (black) SICF curves for the ADM (left) and FDI (right) muscle. **B**) SICF-peak interstimulus intervals. **C**) Individual maximum facilitation for each SICF-peak, relative to test stimulus. *** = P<0.001. Abbreviations: ISI = Interstimulus interval, SICF = short intracortical facilitation, ADM = abductor digiti minimi, FDI = First dorsal interosseous, MEP = motor evoked potential.

### SICF mapping of corticomotor representations of intrinsic hand muscles

SICF mapping caused an anterior-to-posterior shift of the CoG for both intrinsic hand muscles compared to the CoG revealed by single-pulse mapping without changing the medio-lateral distance between the CoGs of the FDI and ADM muscles (Fig. 3A & Fig. 5). Consequently, there were no significant interactions between condition and muscle in any of the models (P>0.39). The magnitude of the anterior-to-posterior shift of the CoGs was positively associated with the peak latency in the SICF_1_ condition (Fig. 3C). The longer the SICF_1_ peak latency, the larger was the anterior-to-posterior shift of the CoGs (P=0.008). This was not the case for the SICF_2_ condition.

**Figure 3.**
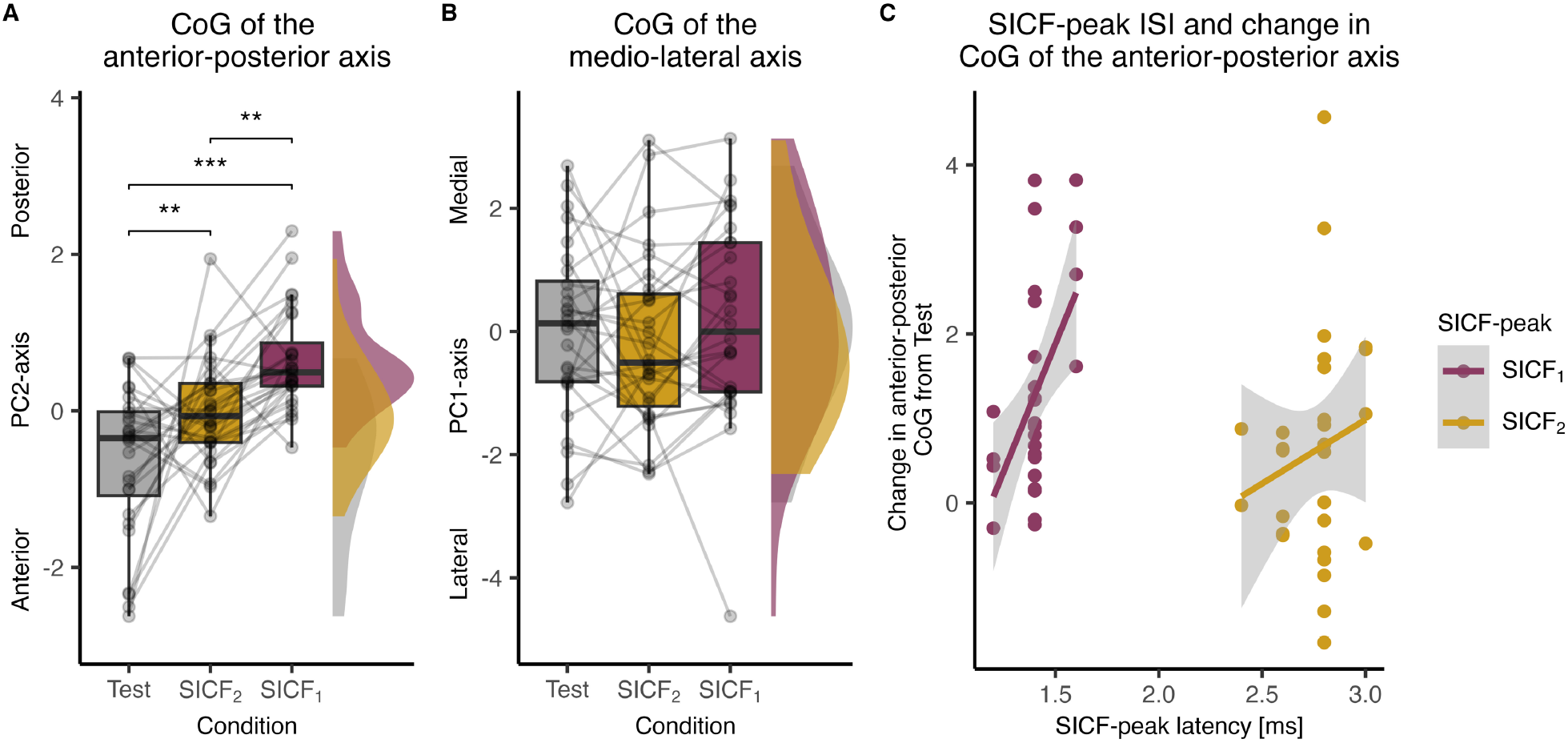
Effects of condition on the center of gravity. Point, box and density plots of within subject mean adjusted center of gravity of the **A**) anterior-posterior axis and **B**) of the medio lateral axis. **C**) Relationship between individual SICF-peak inter stimulus interval and changes in center of gravity of the anterior-posterior axis. ** = P<0.01, *** = P<0.001. Abbreviations: CoG = center of gravity, SICF = short intracortical facilitation, ISI = interstimulus interval.

#### Effects of condition on CoG location

The linear mixed model of PC2 (i.e anterior-posterior axis) CoGs showed a strong effect of *condition* (F_(2)_=18.29, P<0.001). Post-hoc analysis showed that CoGs of both, the FDI and ADM muscles, were located more posterior during SICF_2_ mapping compared to test (beta estimate +/-standard error; 0.62 +/-0.21, P=0.01). Importantly, SICF_1_ CoGs were even more posterior compared to both, the CoGs during single-pulse (1.28 +/-0.21, P<0.001) and SICF_2_ mapping (0.66 +/-0.21, P=0.005). Along the PC1-axis, there was no effect of *stimulation condition* on CoG location (F_(2)_=0.77, P=0.47)(Fig. 3A & B). There were no differences in Euclidian distance between muscles or among the three stimulation conditions (F_(2)_=1.12, P=0.34).

#### Effects of muscle on CoG location

Along the PC1-axis, all three mapping conditions showed medial to lateral separation of CoGs for the ADM and FDI muscles (F_(1)_=30.65, P<0.001) with the FDI muscle positioned more laterally than the ADM in agreement with previous TMS and fMRI mapping studies (15, 16, 29). The CoGs of the FDI and ADM muscles also showed a non-significant difference along the PC2 axis (F_(1)_=3.68, P=0.08) with the CoG of the FDI muscle being more posteriorly positioned than the CoG of the ADM muscle (Fig. 4). Taken together, the data show that the peak corticomotor representations of FDI and ADM, as indexed by the CoG, shifted in the anterior-to-posterior direction on the precentral gyrus during SICF-mapping relative to single-pulse mapping, while maintaining their relative positions in the anterior-posterior and medio-lateral dimension (Fig. 5).

**Figure 4.**
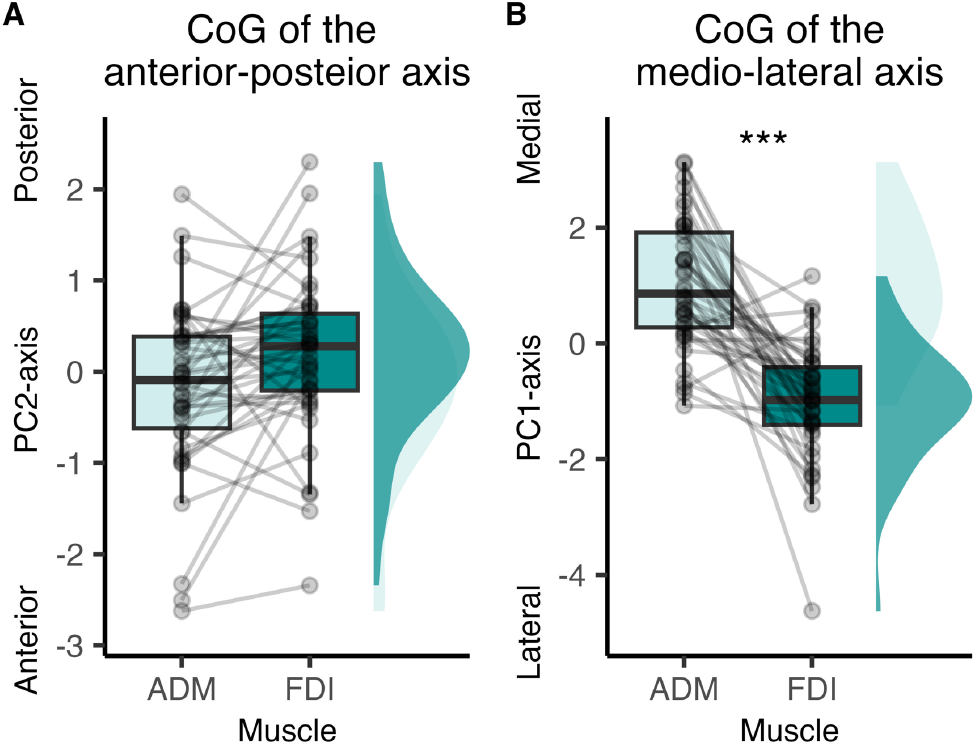
Effects of muscle on center of gravity. Point, box and density plots of within subject mean adjusted center of gravity of the **A**) anterior-posterior axis and **B**) the medio lateral axis. *** = P<0.001. Abbreviations: CoG = center of gravity, ADM = abducter digiti minimi, FDI = first dorsal interosseous.

**Figure 5.**
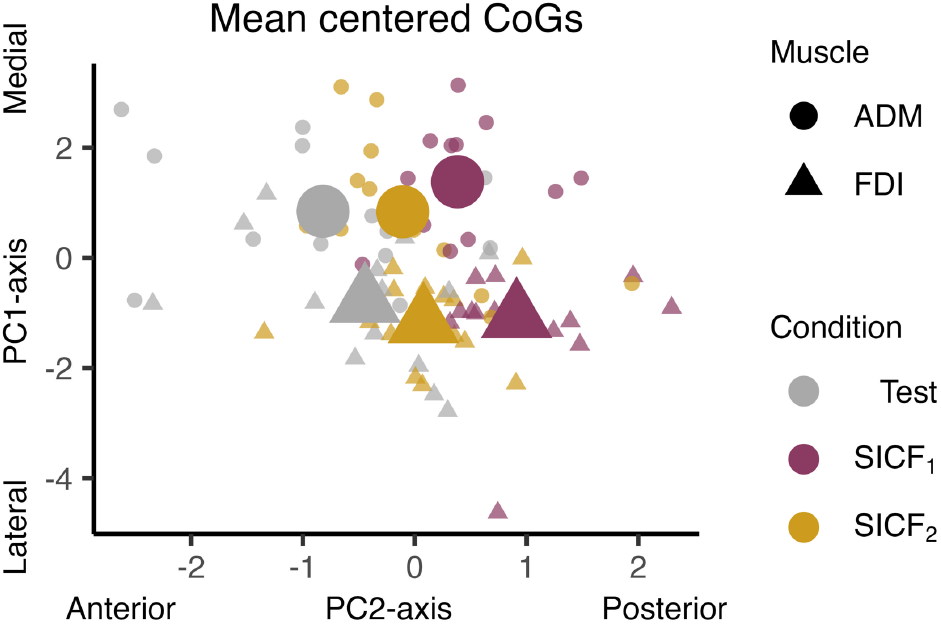
Center of gravity overview. Individual (small) and mean (large), within subject mean adjusted center of gravity in the PC1 and PC2 axes. For visualization purposes only.

#### Sensitivity analyses

To ensure that our results were not biased by the reduction of TMS targets into a plane we performed the same analysis in 3D cartesian space on all three directions and also on a simple 3×7 grid of targets which did not consider the actual placement of TMS targets in relation to each other. These analyses yielded similar results as our primary analysis, showing a separation of the two muscles in all three axes for the 3D map and both axes on the simple grid map (P<0.05). Interestingly, we found that the FDI muscle was more posterior on the grid map, while it was more anterior in 3D cartesian space reflecting that the central sulcus also runs in a posterior to anterior direction. These analyses also showed a separation of the three conditions across the three lines on the simple grid map (P<0.05)(i.e. anterior-posterior direction) and along the actual anterior-posterior direction in 3D cartesian space (P<0.05).

### Relationship between CoG shift and SICF-peak latencies

We conducted mixed linear models with stimulation condition and muscle as fixed effects, subject as a random effect and the difference in CoG along the anterior-posterior PC2-axis as the outcome variable, for each SICF condition separately. These models tested the hypothesis that the spatial shift in CoG position between SICF and single-pulse maps scaled linearly with individual SICF-peak latencies. For SICF_1_ mapping, we found a positive linear relationship between SICF_1_-peak latency and the relative shift in CoG during SICF_1_ mapping (Fig. 3C). The larger the anterior-to-posterior shift in SICF_1_ CoG relative to single-pulse stimulation, the longer was the individual SICF_1_ latency (F(1)=9.45, P=0.008). The SICF_1_ model showed no effect of muscle and no interaction between muscle and stimulation condition (P>0.47). For SICF_2_ mapping, there was no linear relation between SICF_2_-peak latency and the relative shift in CoG during SICF_2_ mapping.

### Effects of SICF mapping on map excitability

As expected, area and volume of both SICF maps were larger than the single-pulse map. Moreover, the volume of the peak normalized SICF maps were larger than the peak-normalized single-pulse map suggesting an expansion of the motor representation independent of the amplitude facilitation.

SICF mapping map area increased from 7.6 +-3.37 (mean +-SD) active targets, to 10.93 +-3.11 for the SICF_2_ (P<0.001) and 13.25 +-2.98 for the SICF_1_ (P<0.001 compared to both test and SICF_2_)(Fig. 6). For log-transformed map volume there was no interaction between *muscle* and *condition* (F_(2)_=1.15, P=0.32), but there was a significant main effect of *muscle* with FDI having a larger map volume than ADM (F_(1)_ = 9.73, P=0.008). We also saw a significant effect of *condition* (F_(2)_=107.15, P<0.001), where SICF_2_ map volume was larger than test volume (1.19 +/-0.11, P<0.001) and SICF_1_ map volume was larger than test (1.48 +/-0.11, P<0.001) and SICF_2_ (0.29 +/-0.11, P=0.021). The results were similar for the peak-normalized maps, i.e. the relative map volume was larger for FDI compared to ADM (F_(1)_=9.8, P=0.007), and the relative map volume was larger for both SICF_1_ and SICF_2_ mapping compared to test (P<0.001), but not between the two conditions (P=0.09). Together, these results show that SICF stimulation results in a relative expansion of the corticomotor representation compared to single-pulse TMS, and this relative expansion was still present after accounting for differences in peak amplitude (i.e., peak corticospinal excitability).

**Figure 6.**
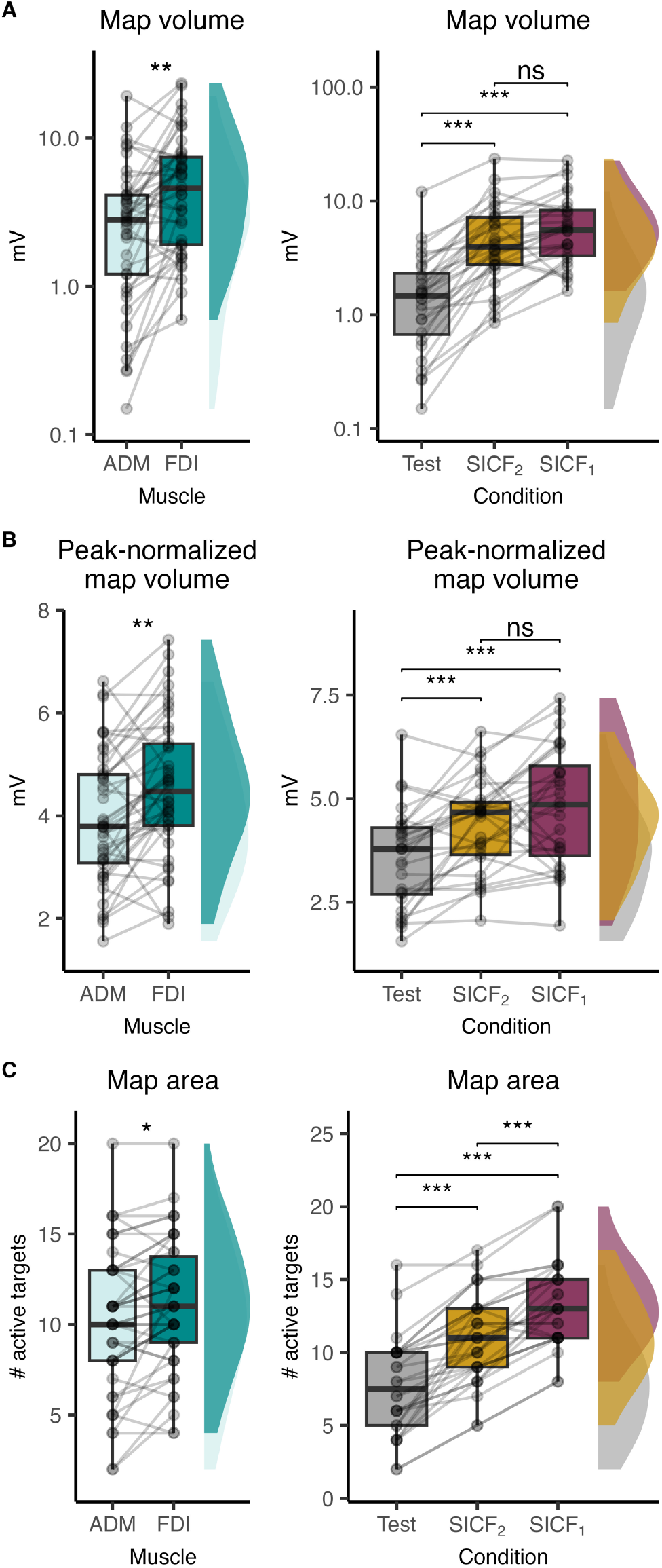
SICF map facilitation. Point, box and density plots of effect of muscle (left) and condition (right) on **A**) map volume, **B**) peak-normalized map volume and **C**) map area. *** = P<0.001.

## Discussion

Sulcus-aligned corticomotor mapping of intrinsic hand muscles revealed that SICF produces a posterior shift in corticomotor representation compared to single-pulse TMS, while preserving the well-known medial-to-lateral somatotopic gradient (16, 30). We also found a spatial discordance in CoGs between SICF_1_ - and SICF_2_ maps, with SICF_1_ CoGs located more posteriorly in the precentral crown. Lastly, exploratory correlational analyses revealed a spatiotemporal relationship between the posterior shift of SICF_1_ CoG and individual SICF_1_ ISI.

### Sulcus-shaped mapping reveal spatially distinct networks mediating SICF

By aligning stimulation targets to the central sulcus and ensuring that the principal current direction is perpendicular to the gyral wall, even small spatial differences between intrinsic hand muscles are detectable (16). This type of sulcus-shaped mapping is highly sensitive to learning dependent plasticity (31) and can be used to probe the intricate organization of the primary motor cortex (15) including inter-individual differences in the anterior-posterior location of single-pulse TMS CoGs (18). We extend these previous findings by showing that sulcus-shaped mapping can reveal spatially distinct corticomotor maps from single- and paired-pulse TMS protocols.

Computational modelling show that TMS primarily activates cortical layers 2-5 in the gyral crown, with layer 5 pyramidal neurons having the lowest threshold followed by layer 2/3 pyramidal interneurons (2). Moreover, at peri-threshold intensities, as applied in this study, direct (D-)waves are typically not observed epidurally (5). Thus, TMS pulses applied in this study therefore likely exerts its excitatory effect on pyramidal tract neurons (PTNs) indirectly through layer 2/3 and 5 pyramidal neurons located in the gyral crown or lip. Early work in animals has demonstrated that I-waves can be triggered from both parietal and premotor sites and the SMA (32-34). Recent evidence also suggests a tight coupling between somatosensory high-frequency oscillations, SICF periodicity and S1 myelination in humans (35). Based on this coupling combined with the notion that TMS directed at the precentral gyrus evokes an (almost) equally strong electrical (e)-field in S1 (1), one hypothesis could be that the posterior shift in SICF hotspot is brought about through co-activation of S1.

Nevertheless, the only factor separating single-pulse and SICF conditions in the current study is the conditioning (second) subthreshold pulse applied in SICF. Since this pulse on its own does not produce any descending activity, the SICF effect must be caused by cortical elements excited by the first supra-threshold pulse. Thus, one interpretation of our results is that the subthreshold conditioning SICF pulse causes supra-threshold excitation of neurons in the subliminal fringe of the first pulse, which are located posterior to neurons excited by low intensity suprathreshold single-pulse TMS.

The relationship between SICF and I-waves is still not known. When two low-intensity pulses with 1-1.4ms, but not 2ms, ISI are applied, the number and size of descending waves recorded epidurally and the ensuing MEP increases (10). However, the use of two supra-threshold stimulations limits the interpretation as the first pulse could be acting on the second pulse rather than the other way around. As such although SICF and corticospinal I-waves likely originate at least in part from the same circuitry, the neural underpinnings are still not known. Recent methodological advancements enable the mapping of immediate TMS evoked EEG activity and may pave the wave for future exploration of this matter (36).

### SICF_1_ and SICF_2_ peaks are generated by spatially distinct networks

Electrophysiological experiments have shown that conditioning SICF with a subthreshold stimulation prior to the test stimuli (i.e. SICI conditioning) produces specific modulation of only the second SICF peak (37). This suggest that distinct inter-neuronal networks mediate the two first SICF peaks. Supporting this notion, only the SICF_2_ peak is modulated during movement preparation (12, 38) and low-level contractions (13) and is facilitated after motor practice (14). We found a systematic spatial difference in SICF_1_ and SICF_2_ CoGs corroborating these previous findings and further suggesting that the two peaks are distinct phenomena generated by two spatially overlapping, but non-identical networks.

Given the previously observed coupling between cortical myelination and SICF periodicity (35), we expected a linear relationship between the temporal distance of SICF peaks and the spatial shift in CoG. Interestingly, we found that the spatial shift of the SICF_2_ peak was smaller than the earlier SICF_1_ peak. This could reflect that SICF_2_ is mediated either by 1) slower conducting unmyelinated interneurons, 2) additional inter-neuronal links, or 3) that the SICF_2_ peak is generated from multiple stimulated sites. Our method is not sensitive enough to dissociate these possibilities, but since we did not find a spatiotemporal relationship between individual SICF_2_ CoG displacement and SICF_2_ peak latency, this could indicate SICF_2_ is influenced by multiple stimulated sites. On the contrary, our exploratory analysis did show a positive relationship between the posterior shift of SICF_1_ CoG and individual SICF_1_-peak ISI. This finding suggests that the longer the projection distance between the SICF_1_ generating network and the hotspot for single-pulse TMS the longer the conduction time. However, further research is needed to support this claim.

### Short-latency paired-pulse TMS facilitates MEP amplitude and expands the corticomotor map

In line with previous studies from preoperative patients, SICF mapping produced both increased map volume and expansion of the map area (39, 40). However, the only study to investigate SICF in neurologically intact humans did not find any change in map area or spatial differences in map CoGs between single-pulse and SICF (41). While the methodological approach was very similar to the current study, the authors used fixed ISIs of 1.4 and 2.8ms and adjusted the TS intensity for SICF to the paired pulse threshold. This rendered both pulses below RMT meaning that it is unlikely that the TS would be sufficiently strong to reveal any spatial differences between the conditions.

Interestingly, our data showed that map expansion was not contingent on the increase in MEP amplitude, but that the spatial extent of intracortical networks underlying SICF exceeds those activated from a single peri-threshold stimulation. Importantly, the map expansion cannot be explained by expansion of the induced e-field from applying two TMS pulses. This has important implications for the use of TMS. First, it shows that the spatial extent of the neural response to TMS in the stimulated cortex is not exclusively determined by the spatial properties of the induced electrical field but dependent on physiological processes triggered by TMS. Second, it can be concluded that single-pulse TMS mapping may not disclose the full extent of the motor eloquent area, which may be relevant for e.g. preoperative TMS mapping (39, 40). More research is needed to dissociate whether map expansion is caused by recruitment of a spatially larger network, a better synchronization of descending corticospinal volleys, or both.

### Methodological considerations

In this study we used biphasic AP-PA TMS pulses which is associated with a low corticomotor threshold (21), enabled us to use lower stimulation intensities compared to monophasic pulses, minimizing the spatial spread of the induced electrical field. Computational modeling of monophasic and biphasic TMS pulses suggest that the posterior lip of the precentral gyrus is more sensitive to the component of the stimulus producing a PA current direction (2). Moreover, monophasic AP and PA oriented pulses may target different I-wave generating networks (42) and potentially also different SICF peaks (43, 44). This could explain the spatial difference between single-pulse and SICF representations, as PA-sensitive neurons in the subliminal fringe could shift the SICF hotspot further posterior. On the other hand, epidural recordings show that I-waves from biphasic AP-PA stimulation occur at the same latency as monophasic PA stimulation (45) and biphasic paired-pulse TMS elicits a characteristic SICF curve (28). These findings suggest that at least at near-threshold intensities, bi- and monophasic TMS target similar networks. Future studies should aim towards SICF mapping using monophasic pulses if possible.

Sulcus-shaped mapping has proven more sensitive to muscle somatotopy than simple grid-based mapping (Raffin et al. 2015), but it does not allow for delineation of the effective cortical site of TMS activation. However, since our mapping procedure was identical between all conditions we are still able to make conclusions about the relative sites of TMS activation. Novel TMS mapping techniques that combines MEP measures with numerical modelling of induced electrical fields have been developed (46-48), which could help elucidate this in future studies.

## Conclusions

Shape-based TMS mapping of precentral motor cortex revealed that single-pulse TMS and SICF engages overlapping, but distinct corticomotor representations of intrinsic hand muscles. This spatial discordance supports the notion that the cortical elements that are most responsive to a single TMS pulse differ from those activated when probing SICF with paired-pulse TMS, revealing yet another facet of the finely arranged organization of the human motor cortex.

